# Spatial aggregation confounds null model-based inference of species association networks

**DOI:** 10.1101/2021.05.25.445651

**Authors:** Andrew J. Rominger

## Abstract

Species association networks inferred from spatially replicated abundance data are typically constructed by comparing observed pairwise similarities to a null distribution generated by permuting the site-by-species matrix while holding row and column marginals fixed. I show that this widely used approach has inflated Type I error whenever individuals within a species are spatially aggregated. High Type I error is driven by the fact that permutational null models do not preserve the shape of the intra-specific spatial species abundance distribution (SSAD). I demonstrate this mechanism with a mathematical argument, simulations of purely neutral communities with no true species associations, and reanalysis of a large, taxonomically and geographically diverse compilation of spatially replicated abundance data. I focus throughout on the specific and widely used class of null-model permutation methods; whether the same artifact affects other approaches to inferring species associations, such as joint species distribution models or the compositional correlation methods common in microbial ecology, remains an open and, I argue, important question for future work.

## Introduction

Species-species and species-environment interactions are fundamental to the evolutionary origins of biodiversity and the maintenance of that diversity in ecological assemblages (Hutchinson, 1961; Holt & Lawton, 1994; Chesson, 2000; Proulx *et al*., 2005; Bascompte & Jordano, 2007; Levine & HilleRisLambers, 2009; Allesina & Tang, 2012; Johnson & Bronstein, 2019). Network theory has come to be a central paradigm in ecology and evolution (May, 1972; Williams & Martinez, 2000; Dunne *et al*., 2002; Proulx *et al*., 2005; Bascompte & Jordano, 2007), and so, not surprisingly, these interactions are often cast as ecological networks where species are nodes (or vertices) and network links (or edges) connect species that interact or share environmental associations.

Abundance or presence/absence data are among the most prevalent and accessible ecological data, and have consequently been frequently leveraged (Morueta-Holme *et al*., 2016; Calatayud *et al*., 2019) in an attempt to infer species interactions and/or associations, beginning at least in the work of Diamond (1975). The specific data used for constructing association networks are often site by species matrices where each cell in the matrix records the abundance (or presence/absence) of a given species at a given site (i.e. sampling location). For simplicity I will refer to both abundance and presence/absence data as simply “abundance data” and focus on true abundance data with the assumed implication that methodological limitations in the case of true abundances will be even more confounding for presence/absence scenarios (Ulrich & Gotelli, 2010).

Two salient insights from decades of debate about using abundance data to infer ecological processes are: 1) interactions in the strict sense cannot be inferred, but *species associations* might be recoverable and these *associations* are assumed to map onto a combination of species-species and species-environment interactions (Morueta-Holme *et al*., 2016; Calatayud *et al*., 2019); and 2) robust null models are necessary to identify which patterns in the data are likely to arise by chance and which are likely to carry biological signal (Ulrich & Gotelli, 2010).

Null models seek to generate plausible data, and patterns in data, while stripping away the actual relationships between species in the data (Ulrich & Gotelli, 2010). Most often a null model is achieved by repeatedly permuting the original data such that any real associations are removed because which species co-occur has been randomized by permutation (Ulrich & Gotelli, 2010). The collection of all such permutations represents a null distribution. These permutation approaches also seek to preserve key aspects of the original data such as maintaining the observed abundances of each species and/or maintaining the observed total number of individuals at each site (Ladau, 2017). These constraints are imposed in an attempt to control the rate of incorrect inference of the presence or absence of network links (Ladau, 2008, 2017). Other methods for inferring association networks do exist, especially model-based inference applied in microbial ecology (e.g. Zhou *et al*., 2010; Kurtz *et al*., 2015), but are less frequently employed in community ecology (but see, e.g., Harris, 2016; Losapio *et al*., 2021).

Here I raise a previously unreported technical issue with the approach of inferring association networks from abundance data using permutational null models. The issue arises from the near-universal spatial aggregation of individuals within species (Legendre, 1993; Condit *et al*., 2000; Lennon, 2000; Dale & Fortin, 2002; McGill & Collins, 2003; Engen *et al*., 2008; Zillio & He, 2010; Harte, 2011; Connolly *et al*., 2017). Using simulation experiments and analysis of real data (from Calatayud *et al*., 2019), I show that intra-specific spatial aggregation (ISSA) causes exceedingly high Type I error rates in permutational null model approaches to inferring association networks, i.e., incorrectly detecting associations when there are none. ISSA in itself is not evidence of species association—it can arise from a large number of processes including simple demographic drift (Kendall, 1949; Hubbell, 2001) in which species do not meaningfully interact with each other or shared environments.

In the absence of ISSA, abundances of a species across sites should follow a Poisson distribution (Diggle, 2003). Thus ISSA is often quantified as the deviation of observed data from a Poisson distribution (Diggle, 2003). Conversely, the negative binomial distribution can model spatially aggregated intra-specific abundances across sites (Bliss & Fisher, 1953; Taylor, 1961; Engen *et al*., 2008; Connolly *et al*., 2017). Critically, the negative binomial arises very naturally from ecological processes as the distribution of population sizes in a birth-death-immigration model (Kendall, 1949). If the abundance of a species at a site is determined by local birth and death with supplemental individuals arriving by immigration from a source pool, then the probability distribution of repeatedly sampled abundances will follow a negative binomial distribution (Kendall, 1949). In this way, simple demographic drift (birth, death, and immigration that are purely stochastic) is expected to produce intra-specific abundances that are negative binomial-distributed and spatially aggregated. Thus ISSA is not, by itself, evidence of species association.

Here I show that spatial aggregation confounds permutational null-model inference because permutation algorithms distort the shape of each species’ spatial abundance distribution (SSAD) toward spatial randomness. Compared to this overly random null, ISSA results in apparent, but spurious, associations.

The technical issue raised here is specific to the widely used family of methods that infer association networks by comparing an observed test statistic to a null distribution generated by Monte Carlo permutation of the site-by-species matrix. I do not test whether the same issue affects methods that do not rely on this style of null model, most notably joint species distribution models, which explicitly condition on measured or latent environmental covariates and site-level random effects (e.g. Zhou *et al*., 2010), or the correlation methods common in microbial ecology (e.g. Kurtz *et al*., 2015). It is not obvious that the mechanism described here transfers directly to these other methods, and I do not want to imply, by omission, that it does. However, any effects of ISSA on joint species distribution modeling and compositional correlation methods should be assessed. Here I focus solely on permutational null models because of their foundational status in the field (Patefield, 1981; Ulrich & Gotelli, 2010) and their continued use in inferring association networks.

## Methods

### Reproducibility and data

A custom R package, *RarePlusComMinus*, accompanies this paper and can be used to reproduce all results. Supplementary Section S1 details how to access and install *RarePlusComMinus*. All analysis scripts and data analyzed are available via the package.

I use a data set of site-by-species abundance matrices compiled by Calatayud *et al*. (2019). The data are included with the *RarePlusComMinus* package as `abundMats` and comprise 385 site-by-species matrices. These compiled data are ideal for investigating the impact of ISSA on association network inference because they were used by Calatayud *et al*. (2019) for the very purpose of network inference and also because they represent a spatially and taxonomically extensive collection of quality-controlled data.

### Quantifying spatial aggregation

I quantify spatial aggregation by characterizing the spatial species abundance distribution (SSAD) for each species in each assemblage. The SSAD is the distribution of a single species’ replicate abundances across sites. I fit both negative binomial and Poisson distributions to the SSADs of each species in each site-by-species matrix using maximum likelihood. The negative binomial distribution is the SSAD that represents ISSA. Specifically, a small dispersion parameter *k* of the negative binomial indicates stronger spatial aggregation. The Poisson is a limiting case of the negative binomial where *k → ∞*.

The `nb_fit()` function from *RarePlusComMinus* fits both negative binomial and Poisson distributions to every species in each site-by-species matrix and returns, for each species from each matrix:

- the negative binomial dispersion parameter (`size`, i.e. *k*) and mean (`mu`)
- the log-likelihoods of negative binomial and Poisson models
- their AIC difference (`delta_AIC`),
- a *χ*^2^-distributed goodness-of-fit statistic for the negative binomial [`zsq`; Etienne (2007)]
- the species’ total abundance and occupancy.

The negative binomial likelihood function does not have a finite maximum when the mean of a species’ abundances across sites is greater than or equal to their variance. A non-finite likelihood is not necessarily evidence against the negative binomial as the true generating distribution; as Supp. Fig. 3 shows, samples drawn from a negative binomial distribution have a nonzero probability of producing an estimated mean greater than or equal to the estimated variance. This occurs disproportionately for rare species and for studies with few sites (Supp. Fig. 2), which is not unexpected: rare species and/or matrices with less sampling will have poorly behaved likelihood functions.

I therefore assess the prevalence of the negative binomial SSAD, and thus ISSA, in two complementary ways:

1. where the likelihood is defined, I compare AIC support for the negative binomial versus the Poisson; and
2. for all species, regardless of whether the likelihood is defined, I evaluate goodness of fit of the negative binomial using the square-transformed *z* statistic of Etienne (2007), which is *χ*^2^-distributed with one degree of freedom under the null hypothesis that the data are negative binomial (Rominger & Merow).

I also fit a log-log linear model relating each species’ dispersion parameter *k* to its relative abundance, which I later use to simulate realistic SSAD shapes for species of any given abundance.

### Quantifying species abundance distributions

I characterize the inter-specific species abundance distribution (SAD, i.e. the distribution of total abundance across species) of each assemblage by fitting three common SAD models and recording the best-supported model and its parameter(s) for each assemblage. The models I consider are the log-series, Poisson log-normal, and zero-truncated negative binomial. I use the `fitSAD()` function from the *pika* package (see Supplementary Section S1 and Supplementary Section S4) to fit model parameters and perform model selection among the three SAD models. SAD model fits will be used in subsequent simulations.

### Quantifying observed and simulated association networks

To construct association networks I calculate pairwise Schoener similarity between each species pair in a site-by-species matrix and compare the observed similarities to those computed from 999 null permuted matrices. I use a fixed-fixed permutation algorithm that preserves row and column marginals. These calculations are carried out by `net_stats()` in the *RarePlusComMinus* package which uses a custom `schoener()` function for similarities and `vegan::nullmodel(x, “r2dtable”)` for null permutations. Each species pair is classified as significantly positively associated, significantly negatively associated, or unassociated at a chosen significance level *α* (default 0.05). I calculate summary statistics of the resulting positive and negative sub-networks (number of vertices and edges, the correlation between abundance and network centrality, and mean and centrality-weighted mean relative abundance), as well as whole-network statistics (connectance, clustering coefficient, average path length, modularity, and the Shannon diversity and mean of the degree distribution).

To evaluate whether estimated network patterns observed in the real data can be produced by spatial aggregation alone, with no true species associations, I simulate site-by-species matrices where every species is truly independent from one another. I simulate both the scenario where there is ISSA (with a negative binomial SSAD) and where there is no ISSA (with a Poisson SSAD). These random site-by-species matrices are simulated as follows:

1. The number of species *S*, number of sites *M*, and shape of the best fitting SAD are sampled (with replacement) from the observed data
2. *S* species abundances *x*_*i*_ … *x*_*S*_ are sampled from the SAD
3. For each species *i* with abundance *x*_*i*_, within-species counts are distributed across the *M* sites according to an SSAD that is either negative binomial (in the case of spatial aggregation) or Poisson (in the case of spatial randomness)
4. The mean of the SSAD is taken to be *x*_*i*_*/M* and, in the case of the negative binomial, the *k* parameter is sampled from a linear model of *k* predicted by relative abundance (Supp. Fig. 5)
5. `net_stats()` is used on the simulated matrix to calculate the same network summary statistics as were calculated for the real data.

*N* = 385 random matrices, the same number as real matrices, are simulated for both the negative binomial and Poisson SSADs.

### Behavior of negative binomial and Poisson SSADs under permutation

To illuminate how negative binomial SSADs behave differently from Poisson SSADs under permutation I simulate many random site-by-species matrices of each type—negative binomial or Poisson. For all SSADs I calculate two summary statistics: the number of occupied sites, and the maximum likelihood estimate of *k* (which serves as a measure of spatial aggregation). I then subject each site by species matrix to permutation and re-calculate the summary statistics. Finally, I compare real and permuted summary statistics to see how permutation has systematically altered their values.

### Mathematical Exploration of Negative Binomial SSADs

To understand what properties of the negative binomial distribution lead to these results I calculate probabilities of different SSAD configurations that would maximize or minimize Schoener similarity.

## Results and Discussion

### Prevalence of negative binomial spatial aggregation

Figure 1 shows that the negative binomial model is strongly preferred over the Poisson for the large majority of species in the Calatayud *et al*. compilation; even where AIC favors the Poisson, it is never strongly favored (Supplementary Section S3). Essentially every case in which the Poisson is favored corresponds to a species for which the negative binomial likelihood is undefined (mean *≥* variance) and sample size is very small (Supp. Fig. 3), consistent with a sampling artifact of rare species rather than genuine evidence for spatial randomness. Indeed, a *z*^2^ goodness of fit test never rejects the negative binomial distribution, even for cases where mean *≥* variance (Supp. Fig. 5).

**Figure 1:**
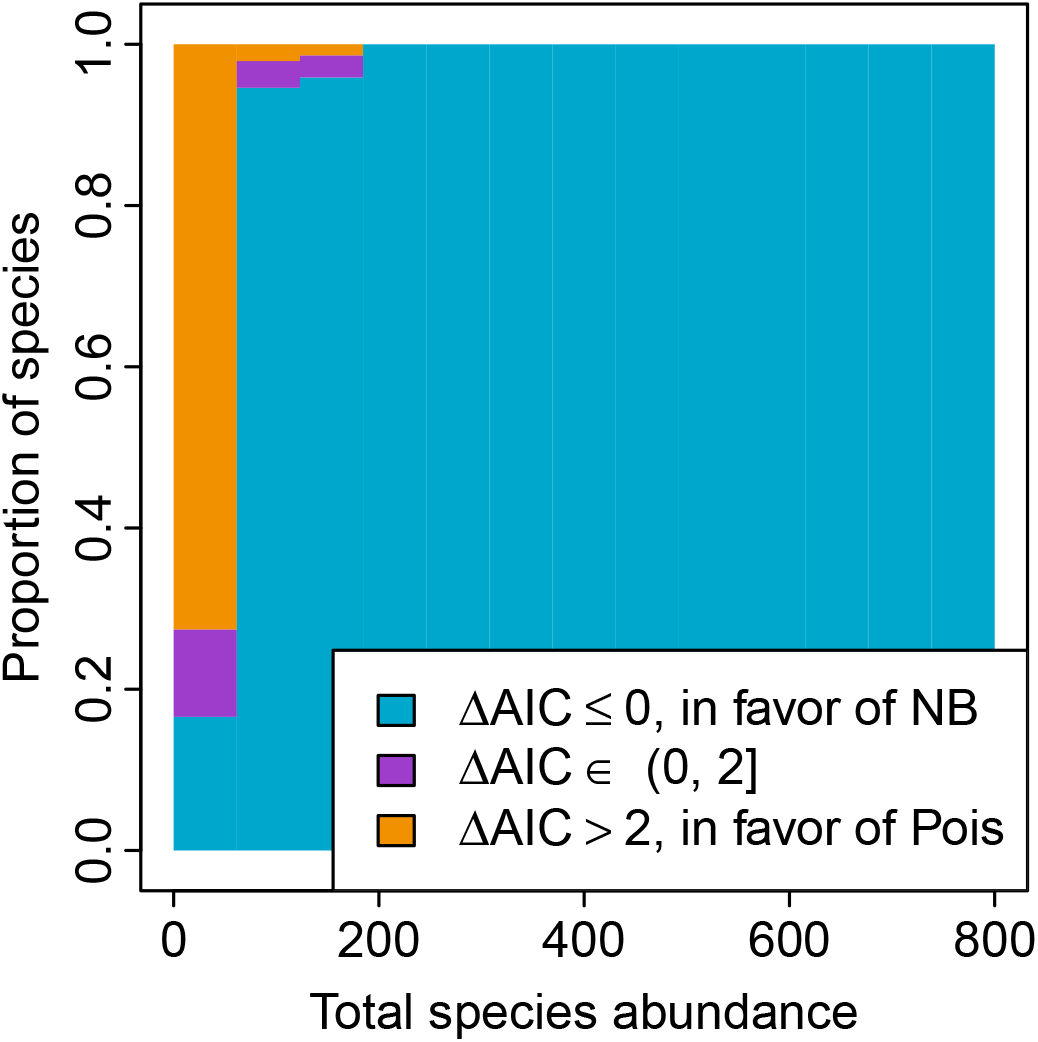
Change in model support for the negative binomial versus the Poisson spatial species abundance distribution (SSAD) as a function of total species abundance. Bars of different colors represent the proportion of species assigned to each level of model support as indicated by the legend.

### Observed and simulated networks

In Figure 2 I first reproduce key results from Calatayud *et al*. (2019)’s Figure 2(B-C). Then to evaluate whether these results can be produced simply from spatial clustering alone I simulate purely random data (with absolutely no association or interaction between species) that match the unevenness of abundances found in the observed data.

**Figure 2:**
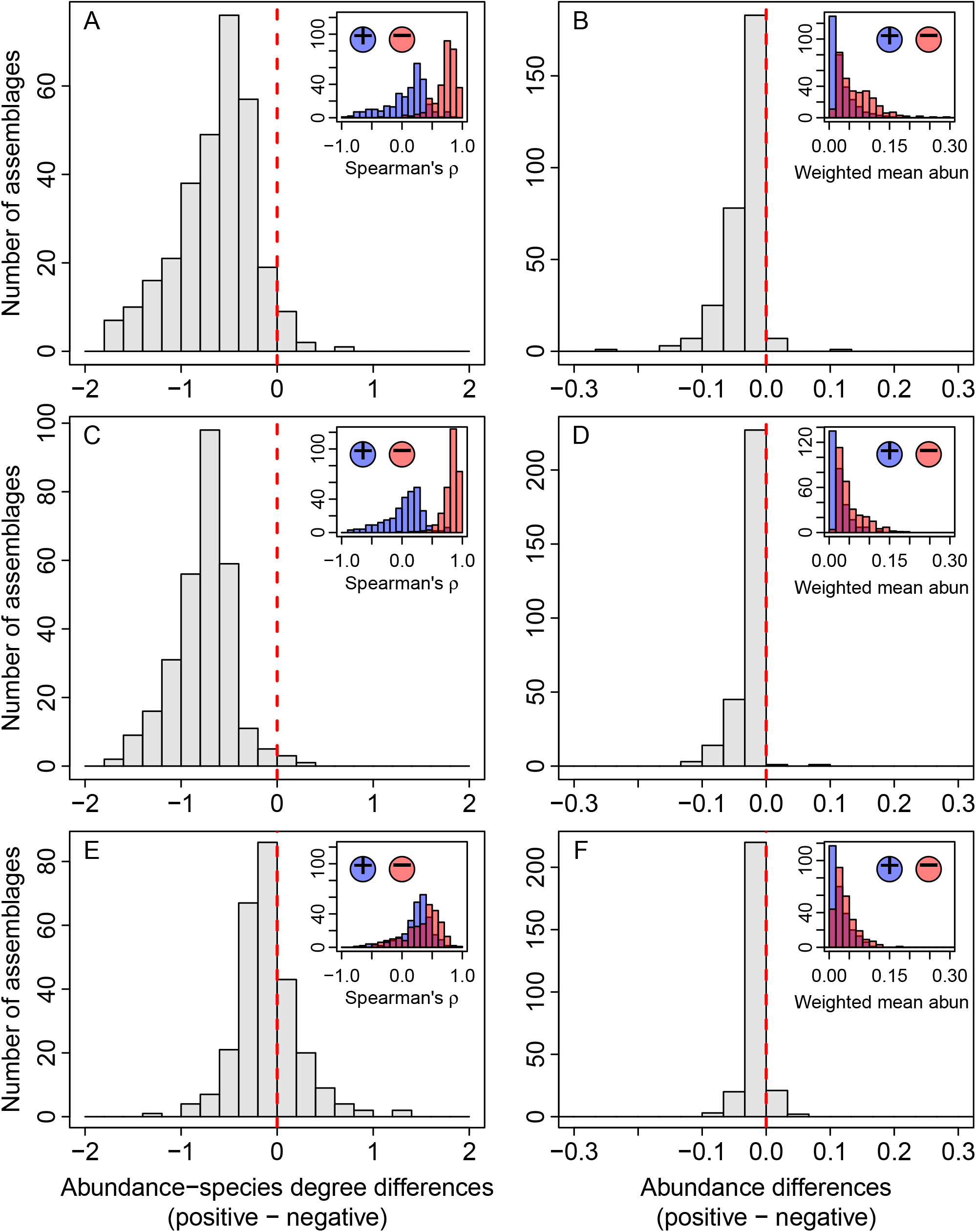
Distributions of correlations between network centrality (i.e. species degree) and abundance (left panels) and distributions of weighted mean abundances (right panels). The main figures show the differences between positive and negative association networks, while the inset figures show the separate distributions for each network. The results of @calatayud2019 Figure 2(B-C) are reproduced here in panels A-B; panels C-D show data simulated with a negative binomial SSAD and no species associations; panels E-F show data simulated with a Poisson SSAD and no species associations.

In the case of a Poisson SSAD the one parameter (the mean) is fully specified by the average site-level abundance of a given species. In the case of a negative binomial SSAD, the mean parameter is again specified by the site-level average, but the size or clustering parameter *k* is not fully specified. To capture the rough features of the data, I sample *k* from a linear relationship (with noise) between the maximum likelihood estimates of *k* and the relative abundance of each species (Supp. Fig. 5).

Figure 2 A–D shows that with a negative binomial SSAD, simulated data closely match observed findings: the correlation between abundance and species’ network degree skews more negative in positive association networks (i.e. more rare species are more highly connected in positive association networks), and positive association networks tend to contain more rare species than negative networks. This correspondence between real and simulated patterns largely disappears when I instead use a Poisson SSAD, highlighting the importance of spatial aggregation in driving the spurious results.

My findings do not depend on simulating SAD and SSAD shapes from the data: in Supp. Fig. 7 I show that the spurious relationship between abundance and association type occurs even when simulating data from just one arbitrary SAD function with the one arbitrary spatially clustered SSAD for all species. In this simulation, again, replacing the spatially clustered SSAD with a Poisson SSAD breaks the spurious connection between abundance and association type as in Figure 2 (E-F).

### Behavior of negative binomial and Poisson SSADs under permutation

Why do negative binomial SSADs reproduce the results while Poisson SSADs fail to? The null model algorithm used here fixes row and column marginals, thus the empirical shape of the SAD is preserved, and the *total* abundances (across all species) at each site are also preserved. However, the way a species’ total abundance is allocated across sites by the null model has a potentially large combinatorial space to explore. In Figure 3 I compare summary statistics of known SSADs to their permuted counterparts and find that the null model transforms negative binomial SSADs to a more Poisson shape, while leaving Poisson SSADs probabilistically unchanged. Specifically, when starting with a negative binomial SSAD, the null model inflates the number of sites individuals are allocated to (more similarly to a Poisson SSAD) and increases the inferred *k* parameter, indicating less spatial clustering in the permuted matrices compared to their non-permuted, negative binomial starting points.

**Figure 3:**
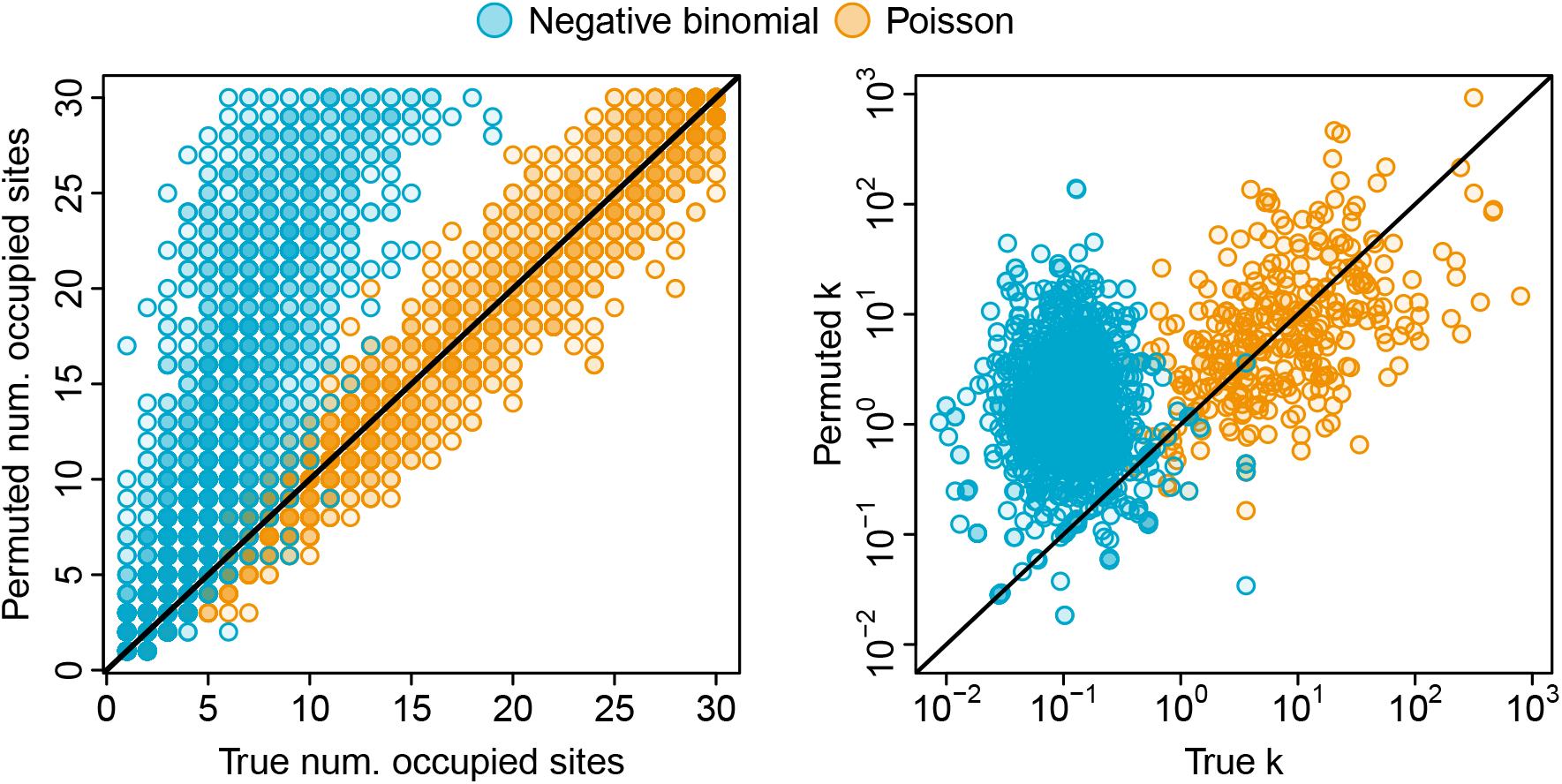
Comparison of SSAD statistics for true and permuted site by species matrices. Colors correspond to the true, un-permuted SSAD. Panel (A) shows how permutation affects number of occupied sites. Panel (B) shows how permutation affects maximum likelihood estimates of the clustering parameter *k*. Points are semi-transparent to help display density. Lines are 1:1 lines.

### Mathematics of negative binomial SSADs

At a mathematical level, clustered SSADs, as compared to spatially even SSADs, increase the probability that rare species will appear positively associated with each other and common species will appear negatively associated. Consider, for example, two rare species: one with a single individual and the other with abundance 5, distributed across 5 sites. Their Schoener similarity is maximized when all individuals occur at the same site, such as this site by species matrix

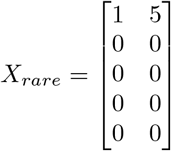

If I define *Q*(*x*_*i*_; *µ* = 1) as the probability of observing *x*_*i*_ individuals in site *i* given an SSAD with mean parameter *µ*, then the probability of the above configuration is *P* (*X*_*rare*_) = *Q*(5; *µ* = 1) *Q*(0; *µ* = 1)^4^. Under a negative binomial SSAD with *k* = 0.1, *P* (*X*_*rare*_) = 4.58 *×* 10^*−*3^ whereas under a Poisson SSAD *P* (*X*_*rare*_) = 5.61 *×* 10^*−*5^.

Conversely, for two common species, say each with abundance 50, an example configuration that *minimizes* their Schoener similarity would be

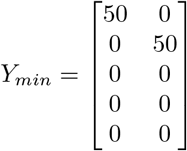

I calculate the probability of any such scenario where no abundances overlap as *P* (*Y*_*min*_) = 4 ((*Q*(50; *µ* = 10)*Q*(0; *µ* = 10)^4^)^2^). With a negative binomial SSAD with *k* = 0.1, *P* (*Y*_*min*_) = 1.41 *×* 10^*−*7^ whereas with a Poisson SSAD *P* (*Y*_*min*_) = 1.61 *×* 10^*−*72^.

I contrast this with a configuration that would *maximize* the Schoener similarity between these two common

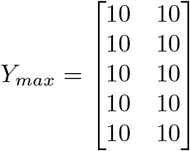

The probability of this configuration is *P* (*Y*_*max*_) = *Q*(10; *µ* = 10)^10^. For the same negative binomial *P* (*Y*_*max*_) = 5.76 *×* 10^*−*22^, and for the Poisson *P* (*Y*_*max*_) = 9.40 *×* 10^*−*10^.

Thus a spatially clustered SSAD, compared to a spatially even SSAD, gives more probability to configurations where rare species appear aggregated and common species appear over-dispersed. Because the null model algorithm permutes site by species matrices to resemble more Poisson-like SSADs, this probabilistic difference between spatially clustered versus even SSADs accounts for the prevalence of rare species in positive association networks and common species in negative association networks.

## Conclusions

Caution should be used when inferring species association from abundance data. More fundamentally than the spurious correlation of abundance with association type, my analysis shows that statistically significant species associations are inferred from data simulated without any real species associations. In data simulated with a negative binomial SSAD, on average 77% of species were placed in positive association networks and 80% in negative association networks with a significance cutoff of *α* = 0.05. With the Poisson SSAD these simulated numbers were 73% for positive networks and 33% for negative networks. For the observed data, on average 74% of species were placed in positive association networks and 67% in negative association networks.

It is becoming increasingly appreciated that abundance data alone are not sufficient to distinguish between different ecological processes (McGill *et al*., 2007; Morlon *et al*., 2009). The question of why rare species persist is fascinating, and Calatayud *et al*. (2019) should be commended for making a concerted effort to illuminate possible mechanisms underlying the phenomenon; however, to reach robust conclusions, other types of data, such as actual experimental measurement of shared environmental associations or species-species interaction strengths, are needed in addition to abundance data.

## Data and Code Availability

All data and code needed to reproduce the results of this manuscript are available at https://github.com/ajrominger/RarePlusComMinus and a detailed description of the analytical approach is available in the supplement.

