## Supplement for "Spatial aggregation confounds null model-based inference of species association networks"

This supplement combines a narrative account of how to reproduce the results in the main text with additional analyses in support of the conclusions in the main text.

#### S1 Reproducibility

The results of this study can be fully reproduced by installing the package *RarePlusComMinus* (available on GitHub) written in R (R Core Team, 2018). Installation requires the package *devtools* (Wickham *et al.*, 2018b), and two other custom packages: *socorro* (Rominger, 2016b) for plotting and *pika* (Rominger, 2016a):

```
dt <- require(devtools)
if(!dt) {
    install.packages('devtools')
    library(devtools)
}

socorroLoad <- require(socorro)
if(!socorroLoad) install_github('ajrominger/socorro')

pikaLoad <- require(pika)
if(!pikaLoad) install_github('ajrominger/pika')

thisPack <- require(RarePlusComMinus)
if(!thisPack) install_github('ajrominger/RarePlusComMinus')
```

All other required packages (Csardi & Nepusz, 2006; Zhang, 2016; Garnier, 2018; Oksanen *et al.*, 2019) are installed with the installation of *RarePlusComMinus*. The *RarePlusComMinus* package includes documented (Wickham *et al.*, 2018a) and unit-tested (Wickham, 2011) functions to carry out all analyses. The help documentation explains these functions, for example

```
?plusMinus
?schoener
```

Now I set up the analysis.

```
library(RarePlusComMinus)
library(pika)
library(socorro)
library(parallel)
library(viridis)

# we can now set caching to be TRUE by default
knitr::opts_chunk$set(cache = TRUE)
```

```
# threading defaults
nthrd <- detectCores()
nthrd <- ifelse(round(nthrd * 0.8) >= nthrd - 1, nthrd - 1, round(nthrd * 0.8))
```

```r
if(nthrd < 1) nthrd <- 1
```

```r
# plotting defaults
parArgs <- list(mar = c(3, 3, 0, 0) + 0.5, mgp = c(1.5, 0.30, 0), tcl = -0.25)
cexDefault <- 1.4
lwdDefault <- 2
figW <- 3.75
figH <- 3.75
```

```r
knitr::opts_chunk$set(fig.width = figW, fig.height = figH, fig.align = 'center')
```

### S2 Reproducing the results of Calatayud *et al.*

First I reproduce some of the key results of Calatayud *et al.* (CEA) (Calatayud *et al.*, 2019), namely how abundances relate to positive and negative association networks. To do this I first process the data from CEA, which I include as `data` in the *RarePlusComMinus* package; more information about the data can be accessed through the R help document via `?abundMats`.

```r
# load data from paper
data('abundMats')

# clean it
obsdat <- lapply(abundMats, function(x) {
    x <- ceiling(x)
    x <- x[rowSums(x) > 0, colSums(x) > 0]

    return(x)
})
```

Now I use the `plusMinus` function to calculate positive and negative association networks and abundances from the observed data. Internally, this function produces `B` (default `B = 999`) randomly permuted matrices using the `r2dtable` algorithm (Patefield, 1981) that fixes row and column marginals. It then compares the Schoener similarities from the original and permuted matrices. I use the custom function `schoener` (made to be more efficient) to compute these similarities, and this function is unit tested against `spaa::niche.overlap`.

```r
commStats <- mclapply(obsdat, mc.cores = nthrd,
                      FUN = function(x) unlist(plusMinus(x)))

commStats <- as.data.frame(do.call(rbind, commStats))

# remove studies with too few plus or minus links
commStats[is.na(commStats$pos.rho.rho) | is.na(commStats$neg.rho.rho), ] <- NA
```

The `plusMinus` function is based on, but not copied from, the function used by CEA in their analyses and made available by the authors at https://figshare.com/articles/Positive_associations_among_rare_species_and_their_persistence_in_ecological_assemblages/9906092. The new `plusMinus` function is streamlined to be faster and thus able to be applied to many more simulations. It is unit tested against CEA's original function.

It should be noted that CEA retained 326 studies after filtering, whereas I retain 305. This is because I removed any dataset for which there were fewer than three links in either the positive or negative networks, whereas CEA removed only those datasets with fewer than two links (Calatayud *et al.*, 2019).

I use these calculations to make Supp. Fig. 1, which reproduces Figure 2 (B-C) from CEA.

```r
# ----
# helper function for making fancy histograms
specialHist <- function(x, breaks, col, add = FALSE, ...) {
    h <- hist(x, breaks, plot = FALSE)

    if(!add) plot(range(h$breaks), range(h$counts), type = 'n', ...)

    rect(xleft = h$breaks[-length(h$breaks)], xright = h$breaks[-1],
         ybottom = 0, ytop = h$counts, col = col)
}

# ----
# helper function for labeling subfigures
figLetter <- function(l, lab, bg = 'transparent', cex = 1, ...) {
    legend('topleft', legend = lab, bg = bg, box.col = 'transparent',
           x.intersp = 0, y.intersp = 0.25, adj = c(0.5, 0.5), cex = cex, ...)
}


# ----
# function to remake Fig 2(b-c)
fig2bc <- function(x, breaksRho, breaksWM, addxlab = TRUE, figLabs = LETTERS[1:2],
                   insetxprop = 0.5, insetyprop = 0.5) {

    # ----
    # relative bounds for inset figures
    insetxMax <- 0.975
    insetxMin <- insetxMax - insetxprop

    insetyMax <- 0.975
    insetyMin <- insetyMax - insetyprop

    # ----
    # set up plot
    plot.new()
    par(parArgs)
    par(cex = 1)

    # ----
    # split into two main plots and fill in
    fi <- split.screen(c(1, 2), erase = FALSE)

    # correlation differences
    screen(fi[1], new = FALSE)

    par(parArgs)
    if(addxlab) {
        par(mar = parArgs$mar + c(0, 0, 0, -0.5))
        xlab <- 'Abundance-species degree differences\n(positive - negative)'
    } else {
        par(mar = parArgs$mar + c(-2.25, 0, 0, -0.5))
        xlab <- ''
    }
```

```
specialHist(x$pos.rho.rho - x$neg.rho.rho, xlim = c(-2, 2),
            breaks = breaksRho, col = 'gray90',
            xlab = '', ylab = 'Number of assemblages')
mtext(xlab, side = 1, line = 2.5)
abline(v = 0, col = 'red', lty = 2, lwd = 2)
figLetter('topleft', figLabs[1])

# raw correlations
fj <- split.screen(matrix(c(insetxMin + 0.02, insetxMax + 0.02,
                             insetyMin, insetyMax), nrow = 1),
                    erase = FALSE)
screen(fj, new = FALSE)

par(parArgs)
par(cex = 0.75)
par(mgp = par('mgp') * par('cex'))

rhoYmax <- 90
rhoFmax <- max(hist(x$pos.rho.rho, breaks = breaksRho / 2, plot = FALSE)$counts,
               hist(x$neg.rho.rho, breaks = breaksRho / 2, plot = FALSE)$counts)

if(0.75 * rhoYmax < rhoFmax + 0.1 * rhoYmax) {
    rhoYmax <- 1.35 * rhoYmax
}

specialHist(x$pos.rho.rho, xlim = c(-1, 1), ylim = c(0, rhoYmax),
            breaks = breaksRho / 2, col = hsv(0.65, alpha = 0.5),
            xlab = expression("Spearman's"~rho),
            ylab = '')
specialHist(x$neg.rho.rho,
            breaks = breaksRho / 2, col = hsv(0, alpha = 0.5),
            add = TRUE)

usr <- par('usr')

points(usr[1] + c(0.2, 0.5) * diff(usr[1:2]), rep(usr[3] + 0.75 * diff(usr[3:4]), 2),
       cex = 3, bg = hsv(c(0.65, 0), alpha = 0.5), pch = 21)
text(usr[1] + c(0.2, 0.5) * diff(usr[1:2]), rep(usr[3] + 0.75 * diff(usr[3:4]), 2),
     labels = c('+', '-'), cex = 2)


# mean differences
screen(fi[2])
par(parArgs)
if(addxlab) {
    par(mar = parArgs$mar + c(0, -1, 0, +0.5))
    xlab <- 'Abundance differences\n(positive - negative)'
} else {
    par(mar = parArgs$mar + c(-2.25, -1, 0, +0.5))
    xlab <- ''
}

specialHist(x$pos.wm - x$neg.wm, xlim = c(-0.3, 0.3),
```

```r
                breaks = breaksWM, col = 'gray90',
                xlab = '', ylab = '')
    mtext(xlab,
          side = 1, line = 2.5)
    abline(v = 0, col = 'red', lty = 2, lwd = 2)
    figLetter('topleft', figLabs[2])


    # raw means
    fk <- split.screen(matrix(c(insetxMin - 0.04, insetxMax - 0.04,
                                insetyMin, insetyMax), nrow = 1),
                       erase = FALSE)
    screen(fk, new = FALSE)

    par(parArgs)
    par(cex = 0.75)
    par(mgp = par('mgp') * par('cex'))

    specialHist(x$pos.wm, xlim = c(0, 0.3),
                breaks = (breaksWM + 0.3) / 2, col = hsv(0.65, alpha = 0.5),
                xlab = 'Weighted mean abund.',
                ylab = '')
    specialHist(x$neg.wm,
                breaks = (breaksWM + 0.3) / 2, col = hsv(0, alpha = 0.5),
                add = TRUE)
    usr <- par('usr')
    points(usr[1] + (1 - c(0.5, 0.2)) * diff(usr[1:2]),
           rep(usr[3] + 0.75 * diff(usr[3:4]), 2),
           cex = 3, bg = hsv(c(0.65, 0), alpha = 0.5), pch = 21)
    text(usr[1] + (1 - c(0.5, 0.2)) * diff(usr[1:2]),
         rep(usr[3] + 0.75 * diff(usr[3:4]), 2),
         labels = c('+', '-'), cex = 2)

    foo <- close.screen(c(fi, fj, fk))

    invisible(NULL)
}
```

```r
fig2bc(commStats, breaksRho = seq(-2, 2, by = 1/5),
       breaksWM = seq(-0.3, 0.3, by = 0.1/3))
```

```r
foo <- close.screen(all.screens = TRUE)
```

It should be noted that my inset plot of mean weighted abundances (Supp. Fig. 1 B) differs from CEA Figure 2C in that their y-axis ranges from 0 to 220, while mine ranges from 0 to 120. I suspected the original authors mislabeled their axis, because the scale as presented would suggest over 600 assemblages, while the number should be 326. Re-scaling their axis to range from 0 to 120 brings the rough estimate from their figure more in line with the reported number of assemblages.

### S3 Exploring patterns of species abundance across space

Now I explore the shape of the spatial species abundance distributions (SSADs) in the real data provided by CEA (Calatayud *et al.*, 2019). The main function I use is `nbFit` from the *RarePlusComMinus* package which calculates summary statistics about the negative binomial and Poisson fits to SSAD data.

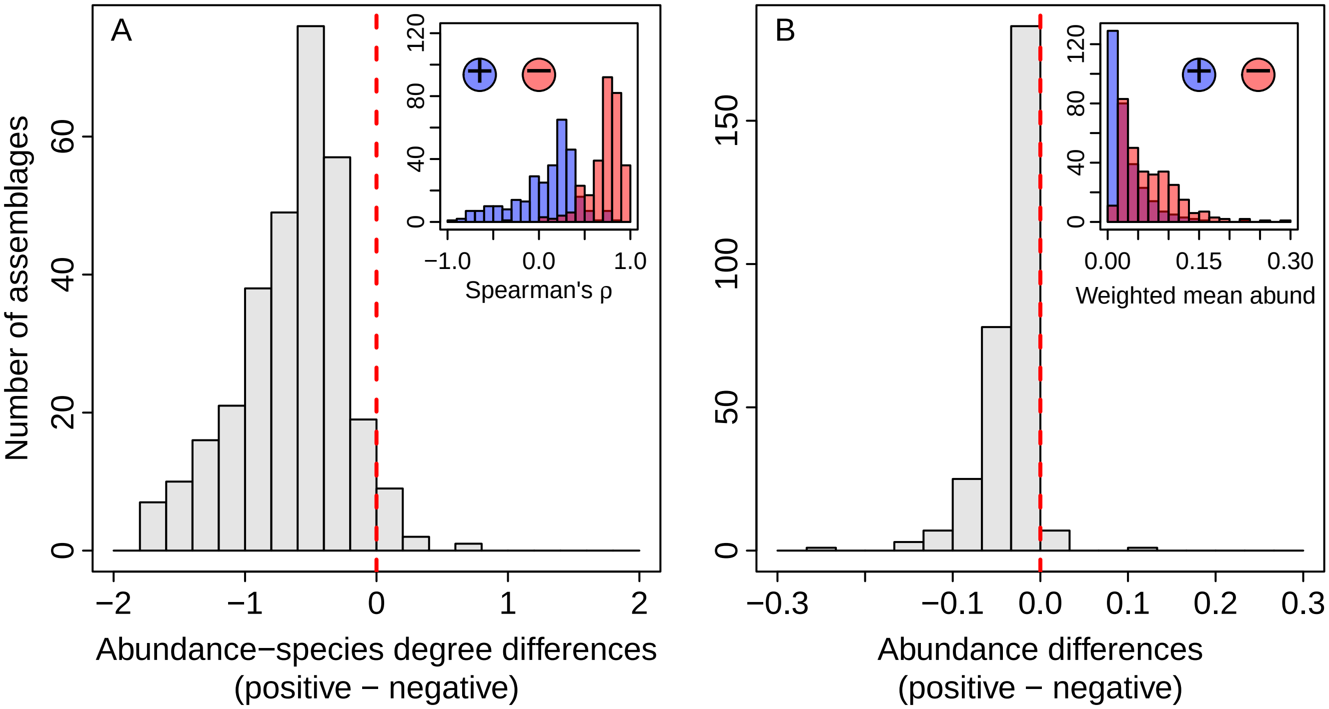


Supplementary Figure 1: Observed relationship between abundance and inferred positive and negative association networks, reproducing Figure 2(B-C) from CEA. This figure is also found in the main text and discussed further there.

```
# limit to only those communities that yielded meaningful networks
summStats <- mclapply(obsdat[rownames(commStats[!is.na(commStats$pos.n), ])],
                      mc.cores = nthrd, FUN = function(x) {
    nbInfo <- nbFit(x)
    cbind(nsite = nrow(x), nspp = ncol(x), J = sum(x), nbInfo)
})

summStats <- data.frame(study = rep(rownames(commStats[!is.na(commStats$pos.n), ]),
                                    sapply(summStats, nrow)),
                        do.call(rbind, summStats))
```

Some care is needed when fitting the negative binomial distribution to data. The negative binomial likelihood function does not have a finite maximum when the mean of the data is greater than or equal to the variance. In the data compiled by CEA I see that the probability of this condition arising is greatest for rare species and also when the number of spatial replicates is small (especially when the number of spatial replicates is equal to 10; Supp. Fig. 2). The fact that numerical issues arise with small sample size and low abundance should not be surprising (these cases are subject to sampling issues), and indeed, even when the generating distribution is a negative binomial, I expect that rare species will often be sampled in such a way as to produce numerical issues (Supp. Fig. 3).

```
abundNsite <- as.matrix(expand.grid(abund = 1:40, nsite = 9:90))

finiteK <- sapply(1:nrow(abundNsite), function(i) {
    j <- summStats$abund >= abundNsite[i, 1] & summStats$nsite >= abundNsite[i, 2]
    N <- sum(j)
    good <- sum(is.finite(summStats$size[j]))
```

```r
    return(good / N)
})
```

```r
colcut <- 0.8
colmin <- min(finiteK)

propLo <- (colcut - colmin) / (1 - colmin)
ncolz <- 100
nLo <- round(ncolz * propLo)
nHi <- ncolz - nLo
colLo <- rev(magma(nLo, begin = 0.3, end = 0.7))
colHi <- viridis(nHi, begin = 0.4)


finiteK <- list(x = sort(unique(abundNsite[, 1])),
                y = sort(unique(abundNsite[, 2])),
                z = matrix(finiteK, nrow = length(unique(abundNsite[, 1])),
                           ncol = length(unique(abundNsite[, 2]))))

# ----
# plotting

layout(matrix(1:2, ncol = 2), widths = c(4, 1))
par(parArgs)

image(finiteK$x, finiteK$y, ifelse(finiteK$z <= colcut, finiteK$z, NA), col = colLo,
      xlab = 'Total species abundance', ylab = 'Number of sites')
image(finiteK$x, finiteK$y, ifelse(finiteK$z > colcut, finiteK$z, NA), col = colHi,
      add = TRUE)

contour(finiteK$x, finiteK$y, finiteK$z,
        method = 'edge', levels = c(1), col = 'white', drawlabels = FALSE, lwd = 3,
        add = TRUE)
box()


par(parArgs)
par(mar = c(parArgs$mar[1], 0.5, parArgs$mar[3], parArgs$mar[2]))

plot(1:2, ylim = c(colmin, 1), type = 'n', axes = FALSE, xlab = '', ylab = '',
     xaxs = 'i', yaxs = 'i')

rect(xleft = 1, xright = 2,
     ybottom = seq(colmin, 1, length.out = ncolz + 1)[-(ncolz + 1)],
     ytop = seq(colmin, 1, length.out = ncolz + 1)[-1],
     pch = 16, col = c(colLo, colHi), border = c(colLo, colHi))
box()

atz <- pretty(c(colmin, 1))
atz <- atz[atz >= colmin]
axis(4, at = atz)

mtext('Proportion of species with solveable MLE', side = 4, line = parArgs$mgp[1])
```

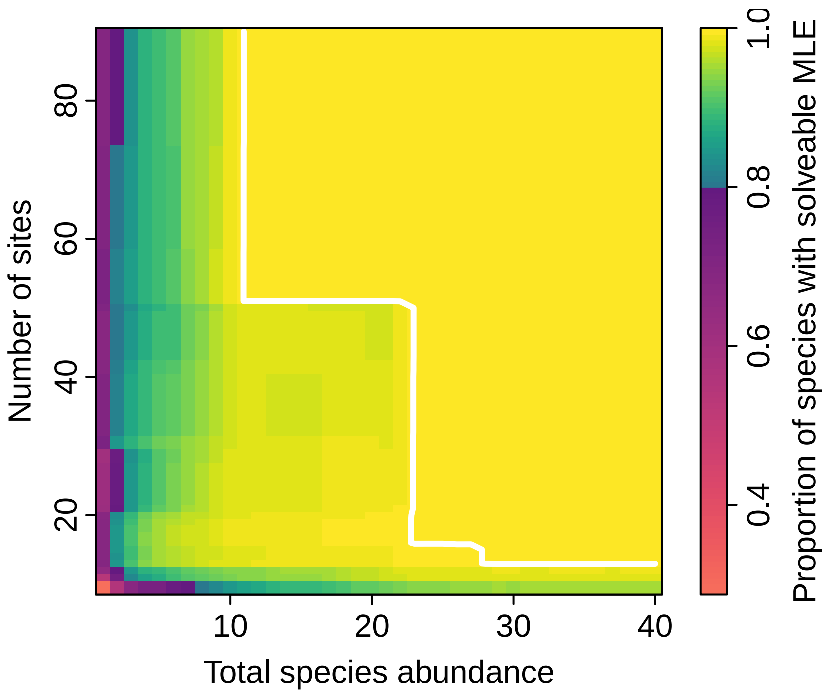


Supplementary Figure 2: Relationship between the solvability of the negative binomial likelihood function and the number of spatial replicates (sites) and total species abundances found in the real data. The color gradient indicates the cumulative proportion of species-level observations in the data for which a solution can be found for the negative binomial likelihood function. By cumulative I mean that the proportion in each cell represents all species with at least that abundance and at least that many spatial replicates. The white line shows the cutoff above which all data yield solvable negative binomial likelihoods.

```r
pmleGood <- function(N, k, nsite) {
    o <- mclapply(N, mc.cores = nthrd, FUN = function(n) {
        o <- replicate(500, {
            x <- rnbinom(nsite, k, mu = n / nsite)
            mean(x) < var(x)
        })

        return(mean(o))
    })

    return(unlist(o))
}

N <- 1:100
hiK <- 1
loK <- 0.1
pgood10loK <- pmleGood(N, loK, 10)
pgood100loK <- pmleGood(N, loK, 100)
pgood10hiK <- pmleGood(N, hiK, 10)
pgood100hiK <- pmleGood(N, hiK, 100)
```

```r
par(parArgs)

plot(N, pgood10loK, type = 'l', ylim = 0:1, lwd = lwdDefault, col = gray(0.6),
     xlab = 'Total species abundance', ylab = 'Probability of solvable NB likelihood')
lines(N, pgood100loK, lwd = lwdDefault)
lines(N, pgood10hiK, lwd = lwdDefault, col = hsv(0, 0.6))
lines(N, pgood100hiK, lwd = lwdDefault, col = hsv(0))

legend('bottomright', legend = paste0('Num site = ', rep(c(10, 100), each = 2), '; ',
                                      'k = ', rep(c(loK, hiK), 2)),
       lty = 1, lwd = lwdDefault, col = hsv(0, c(0, 0, 0.6, 1), c(0.6, 0, 1, 1)),
       bty = 'n')
```

Thus rare species are not very informative about the shape of this distribution (again, this should not be surprising). Rare species are also the most prevalent in natural communities, thus when I look at the real data it seems that the Poisson most often beats the negative binomial in terms of AIC. But when I look at this in terms of abundance I see that the Poisson clearly wins (i.e. $\Delta\text{AIC} > 2$) only for cases when total species abundance is small (Supp. Fig. 4). In cases where $\Delta$AIC favors the Poisson but there is little discrimination between models (i.e. $0 < \Delta\text{AIC} \leq 2$) I still see that abundances tend to be small. When the negative binomial clearly wins (i.e. $\Delta\text{AIC} \leq -2$), it is for higher abundances.

```r
img1D <- function(subx, allx) {
    o <- hist(subx[subx <= 800],
              breaks = seq(0,
                           800,
                           length.out = 14),
              plot = FALSE)

    return(o[c('breaks', 'counts')])
}

poWin <- img1D(summStats$abund[summStats$dAIC > 2], summStats$abund)
mixWin <- img1D(summStats$abund[summStats$dAIC > 0 & summStats$dAIC < 2], summStats$abund)
```

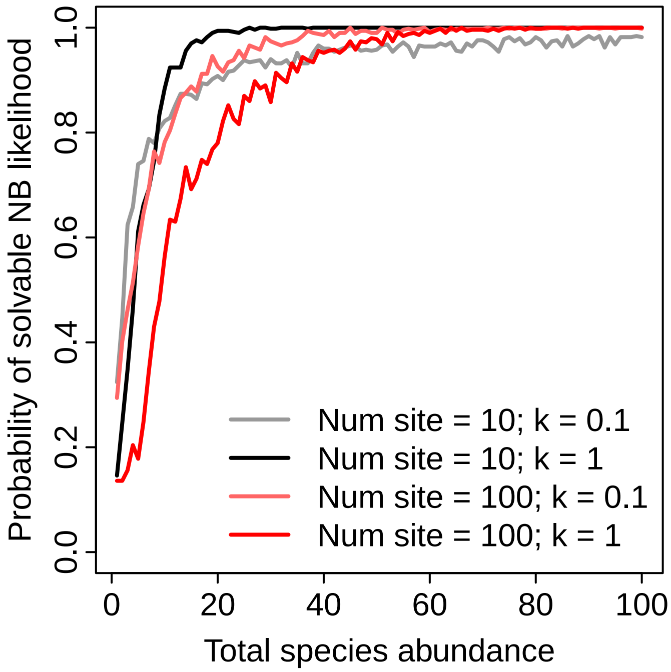


Supplementary Figure 3: The relationship between total species abundance and the probability that the negative binomial likelihood function will be solvable. Different line colors correspond to different negative binomial parameterizations and different number of spatial replicates.

```
nbWin <- img1D(summStats$abund[summStats$dAIC <= 0], summStats$abund)

llImg <- cbind(nbWin$counts, mixWin$counts, poWin$counts)
llImg <- llImg / rowSums(llImg)
llImg[llImg == 0] <- NA


colz <- hsv(c(0.53, 0.78, 0.1), c(1, 0.7, 1), c(0.8, 0.8, 0.95))

par(parArgs)
barplot(t(llImg), space = 0, border = NA, col = colz, ylim = c(-0.05, 1.05),
        xlab = 'Total species abundance', ylab = 'Proportion of species')
box()

axis(1, at = approxfun(nbWin$breaks, 1:length(nbWin$breaks))(pretty(nbWin$breaks)) - 1,
     labels = pretty(nbWin$breaks))

legend('bottomright', legend = expression(Delta*AIC <= 0*', in favor of NB',
                                          Delta*AIC %in%~'(0, 2]',
                                          Delta*AIC > 2*', in favor of Pois'),
       fill = colz,
       bg = 'white')
```

Additionally, the maximum ΔAIC in favor of the Poisson is 2.073904 which is only very slightly above the threshold of 2 often considered to be the cutoff between models with weakly or strongly different support (Burnham & Anderson, 2003). All cases in which ΔAIC favors the Poisson by at least 2 are also those cases in which the mean of the data is greater than or equal to the variance, i.e. where the negative binomial cannot

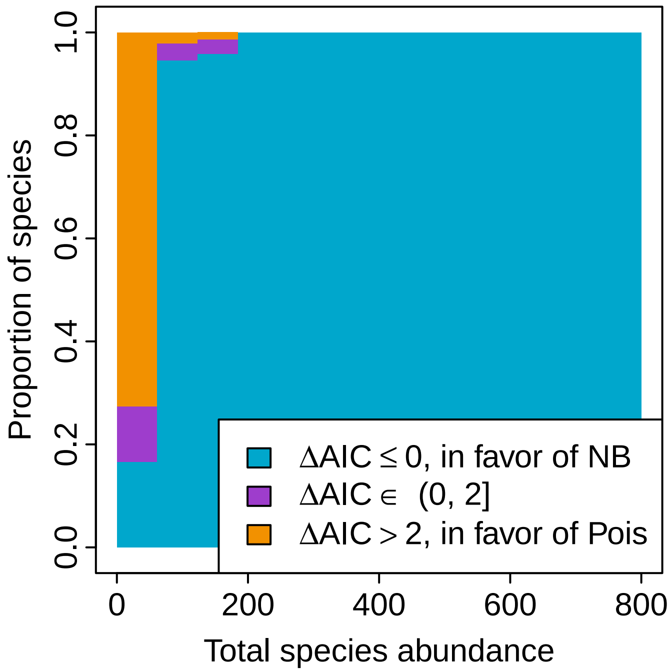


Supplementary Figure 4: Change in model support for the negative binomial versus the Poisson spatial species abundance distribution (SSAD) as a function of total species abundance. Bars of different colors represent the proportion of species assigned to each level of model support as indicated by the legend.

be fit.

Taking all this into consideration, I rely on a different test to demonstrate the good fit of the negative binomial to the data. In Supp. Fig. 5 I show that the data are well fit by the negative binomial via a likelihood-based goodness of fit test (Rominger & Merow; Etienne, 2007). This test scales the observed likelihood by the sampling distribution of likelihoods given the hypothesis that the negative binomial is the correct distribution. This test statistic, when squared, follows a $\chi$-squared distribution with 1 degree of freedom (Rominger & Merow), allowing me to make a parametric cutoff of when the data are not well represented by a negative binomial. No assemblage analyzed rejected the negative binomial distribution.

In Supp. Fig. 5 I also explore the relationship of the clustering parameter $k$ with species relative abundance. The purpose of this latter analysis is to be able to simulate random but realistic data.

```
# data for linear model of k, excluding sites that didn't produce a good network
dat4k <- with(summStats[summStats$study %in%
                            rownames(commStats[!is.na(commStats$pos.n), ]) &
                            is.finite(summStats$size), ],
              data.frame(logk = log(size),
                         loga = log(abund / J),
                         logS = log(nspp))
)


kMod <- lm(logk ~ loga, data = dat4k)

kfun <- function(nspp, abund) {
    J <- sum(abund)

    exp(predict(kMod, newdata = data.frame(loga = log(abund / J))) +
```

```
              rnorm(nspp, sd = summary(kMod)$sigma))
}
```

```
layout(matrix(1:2, nrow = 1))

par(parArgs)
plot(summStats$abund, summStats$z, log = 'xy',
     ylim = range(summStats$z, qchisq(0.999, 1)),
     xaxt = 'n', yaxt = 'n',
     xlab = 'Abundance', ylab = expression('Goodness of fit '~z^2))
logAxis(1:2, expLab = TRUE)
abline(h = qchisq(0.95, 1), col = 'red', lwd = 2)
figLetter('topleft', 'A', bg = 'white')
box()

plot(summStats$abund / summStats$J, summStats$size, log = 'xy', xaxt = 'n', yaxt = 'n',
     xlab = 'Relative abundance', ylab = expression(k))
curve(kMod$coefficients[1] * x^kMod$coefficients[2], col = hsv(0.56, 0.6, 0.8),
      lwd = 2, add = TRUE)
logAxis(1:2, expLab = TRUE)
figLetter('topleft', 'B')
```

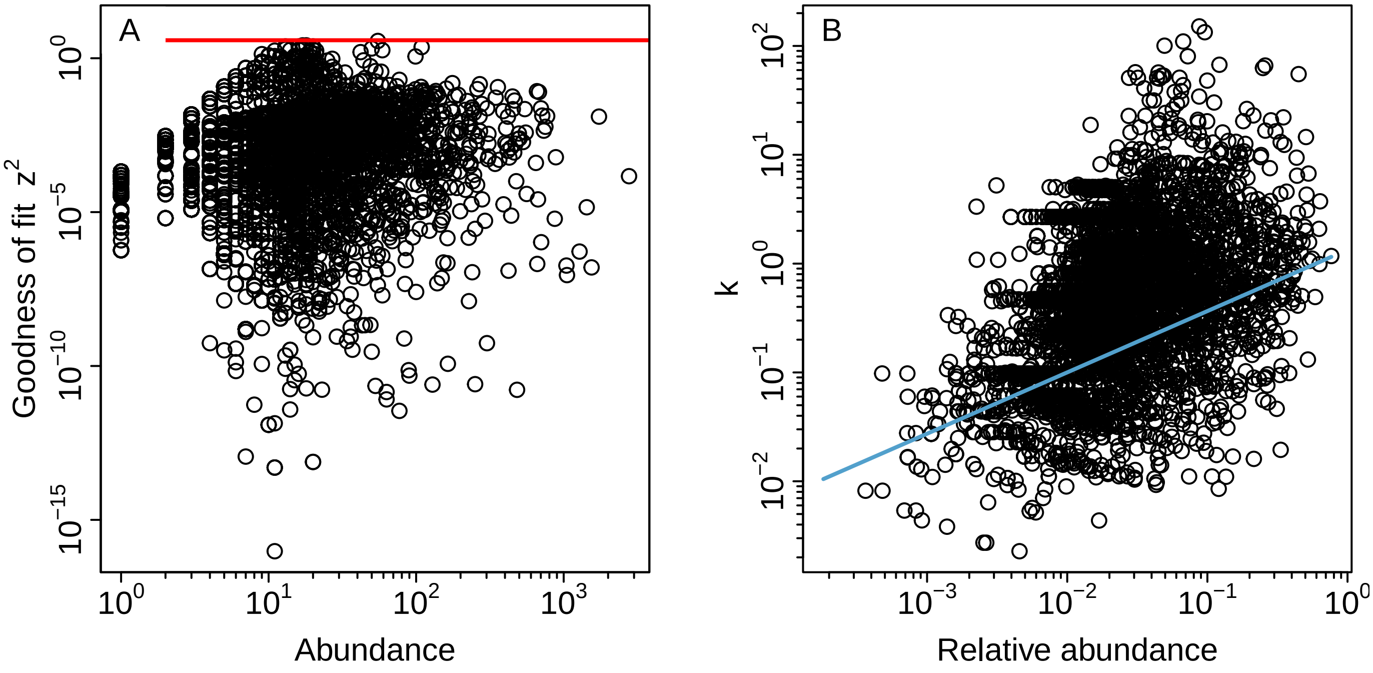


Supplementary Figure 5: Observed spatial species abundance distributions (SSAD) as characterized by (A) the goodness of fit of the negative binomial distribution and (B) the relationship between a given species' relative abundance and its clustering parameter $k$. The horizontal red line in (A) indicates the critical value above which we would reject the negative binomial; no points are above this line. The blue regression line in (B) shows the best fit log-log linear model.

#### S4 Exploring the species abundance distribution

I also need to know the shapes of the species abundance distributions (SAD) of each assemblage to simulate realistic data. I do this using the function `fitSAD` from the custom *pika* package. I perform model selection on three standard SAD forms: the log-series, Poisson log-normal, and zero-truncated negative binomial, and record the best-fit model and its parameter(s) for each assemblage.

```
mods <- c('lseries', 'plnorm', 'tnegb')
sadStats <- mclapply(obsdat, mc.cores = nthrd, FUN = function(x) {
    nsite <- nrow(x)
    x <- colSums(x)
    s <- fitSAD(x, mods)

    i <- which.min(sapply(s, AIC))
    o <- s[[i]]$MLE
    if(i == 1) o <- c(o, NA)

    o <- c(i, o, sum(x), length(x), nsite)
    names(o) <- NULL

    return(o)
})

# mclapply returns 'try-error' objects (length 1) when a worker fails; surface the
# underlying message instead of failing obscurely downstream
sadErr <- sapply(sadStats, function(x) inherits(x, 'try-error'))
if(any(sadErr)) stop('fitSAD failed for ', sum(sadErr), ' assemblage(s); first error: ',
                     sadStats[[which(sadErr)[1]]])

sadStats <- as.data.frame(do.call(rbind, sadStats))
names(sadStats) <- c('mod', 'par1', 'par2', 'J', 'nspp', 'nsite')
sadStats$mod <- mods[sadStats$mod]

# limit to only those sites that produced good networks
sadStats <- sadStats[rownames(sadStats) %in%
                         rownames(commStats[!is.na(commStats$pos.n), ]), ]
```

```
# helper function to make a rank abundance dist for a hypothetical community of
# `S` species given model `m` and parameters `p`

hypRAD <- function(m, p, S) {
    x <- sad(model = m, par = p[!is.na(p)])

    r <- sad2Rank(x, S)

    return(r / sum(r))
}

S <- 100

allRAD <- mclapply(1:nrow(sadStats), mc.cores = nthrd, FUN = function(i) {
    hypRAD(sadStats$mod[i], as.numeric(sadStats[i, c('par1', 'par2')]), S)
})
allRAD <- do.call(rbind, allRAD)
```

```r
radEnv <- lapply(unique(sadStats$mod), function(m) {
    apply(allRAD[sadStats$mod == m, ], 2, quantile, probs = c(0.025, 0.975))
})
```

To demonstrate the marked unevenness of the SADs, in Supp. Fig. 6 I look at the outline of the shapes of all the rank abundance distributions for a hypothetical community of 100 species.

```r
# ----
# helper function to make overlapping polygons more clear
specialPoly <- function(x, y, col) {
    polygon(x, y, col = colAlpha(col, 0.4), border = NA)
    polygon(x, y, border = col)
}


# ----
# plotting

par(parArgs)

plot(1, xlim = c(1, S), ylim = range(unlist(radEnv)), log = 'y', yaxt = 'n', type = 'n',
     xlab = 'Species rank', ylab = 'Relative abundance')
logAxis(2, expLab = TRUE)

polygon(c(1:S, S:1), c(radEnv[[1]][1, ], rev(radEnv[[1]][2, ])),
        col = hsv(0.56, 1, 0.8, 0.4))
polygon(c(1:S, S:1), c(radEnv[[2]][1, ], rev(radEnv[[2]][2, ])),
        col = hsv(0.05, 1, 1, 0.4))
polygon(c(1:S, S:1), c(radEnv[[3]][1, ], rev(radEnv[[3]][2, ])),
        col = hsv(0.75, 1, 0.8, 0.4))
polygon(c(1:S, S:1), c(radEnv[[1]][1, ], rev(radEnv[[1]][2, ])),
        border = hsv(0.56, 1, 0.8))
polygon(c(1:S, S:1), c(radEnv[[2]][1, ], rev(radEnv[[2]][2, ])),
        border = hsv(0.05, 1, 0.8))
polygon(c(1:S, S:1), c(radEnv[[3]][1, ], rev(radEnv[[3]][2, ])),
        border = hsv(0.75, 1, 0.8))

legend('topright',
       legend = c('Log-series', 'Poisson log-norm', 'zero-trunc. nbinom'),
       pch = 22, pt.cex = 2, pt.lwd = 1.5,
       col = hsv(c(0.56, 0.05, 0.75), 1, 0.8),
       pt.bg = hsv(c(0.56, 0.05, 0.75), 1, c(0.8, 1, 0.8), 0.4),
       bty = 'n')
```

#### S5 Simulating random data and artifactual associations

Now I can simulate abundance data matching the overall shapes of the observed SADs and SSADs but with absolutely no real association between species.

```r
# number of simulations to run
nsim <- round(1.25 * sum(!is.na(commStats$pos.wm)))
```

I simulate 381 random assemblages, slightly more than the 305 observed assemblages because some simulated assemblages will be rejected based on the data standards used. In Figure 1 of the main text I plot these

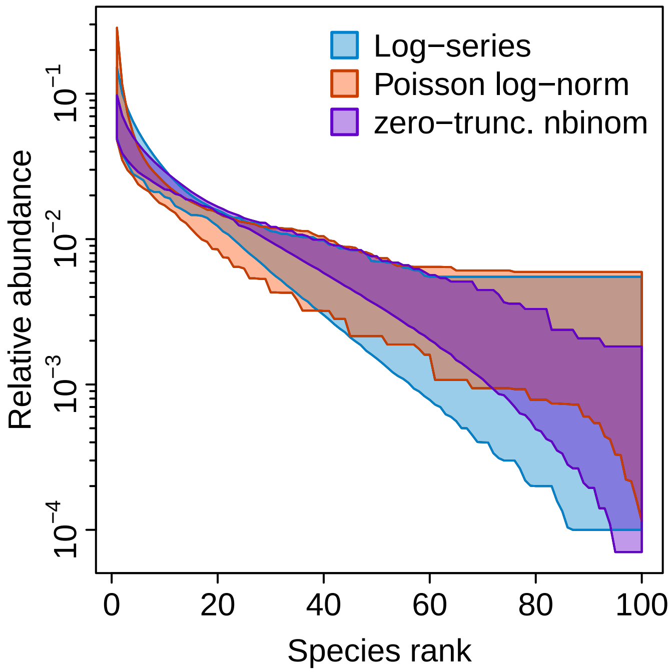


Supplementary Figure 6: Graphical summary of the shapes of the best-fit species abundance distribution models. Polygons represent the 95% confidence envelope of the rank abundance plots for each of the three SAD models considered.

simulated results alongside the results from the real data.

```
# loop over simulation replicates
simPMData <- simPlusMinus(sadStats = sadStats, mcCores = nthrd,
                          ssadType = 'nbinom', kfun = kfun, nsim = nsim)
```

Now I want to figure out if the correspondence between real and simulated results is specifically because of the shape of the SSAD (i.e. negative binomial) or if a spatially unclustered SSAD would also yield so close a match. I do this by modeling the SSAD with a Poisson distribution.

```
# loop over simulation replicates
simPMDataPois <- simPlusMinus(sadStats = sadStats, mcCores = nthrd,
                              ssadType = 'pois', kfun = NULL, nsim = nsim)
```

Lastly, I want to know whether I can produce a spurious relationship between abundance and association type without any reference to the observed data. For this experiment I imagine one arbitrary SAD and combine it with one of two arbitrary SSADs (either negative binomial or Poisson) and see if qualitatively similar results are found.

```
oneK <- 0.1
b <- 0.01
nsiteSimp <- 20
nsppSimp <- 50
nsimSimp <- nsim
```

In this case I use a log-series SAD with $\beta = 0.01$, a negative binomial with $k = $0.1, and consider an assemblage with on average 20 sites and 50 species.

```
simPMSimpNB <- simpleSim(nsiteSimp, nsppSimp, nthrd, sadfun = function(n) rlseries(n, b),
                         ssadfun = function(n, mu) rnbinom(n, oneK, mu = mu),
```

```
                         nsim = nsimSimp)
```

```
simPMSimpPo <- simpleSim(nsiteSimp, nsppSimp, nthrd, sadfun = function(n) rlseries(n, b),
                         ssadfun = function(n, mu) rpois(n, lambda = mu),
                         nsim = nsimSimp)
```

In Supp. Fig. 7 I see that indeed, when I use a realistic SAD and pair it with a negative binomial SSAD, I reconstruct a spurious relationship between rare species and positive associations versus common species and negative associations. When this same SAD shape is paired with a Poisson SSAD, I see this spurious association is substantially reduced.

```
# calculate breaks
wmmax <- ceiling(max(simPMSimpNB$pos.wm, simPMSimpNB$neg.wm,
                     simPMSimpPo$pos.wm, simPMSimpPo$neg.wm,
                     na.rm = TRUE) * 9) / 3

foo <- split.screen(c(2, 1))

screen(1)
fig2bc(simPMSimpNB, breaksRho = seq(-2, 2, by = 1/5),
       breaksWM = seq(-wmmax, wmmax, by = 0.1/3), addxlab = FALSE, figLabs = c('A', 'B'))

screen(2)
fig2bc(simPMSimpPo, breaksRho = seq(-2, 2, by = 1/5),
       breaksWM = seq(-wmmax, wmmax, by = 0.1/3), figLabs = c('C', 'D'))

foo <- close.screen(all.screens = TRUE)
```

```
nsite <- 30
nspp <- 50
nsim <- 100
```

To understand why these spurious results occur I must understand what the fixed-fixed null model (Patefield, 1981; Ulrich & Gotelli, 2010) does to the underlying SSAD. I know that the fixed-fixed algorithm preserves, by definition, the SAD and the total abundances across sites, but within any given species, the allocation of its abundances has a potentially large combinatorial space. To understand what happens to SSADs when permuted, I simulate 100 assemblages, each with 50 species and 30 sites and compare the shape of the permuted SSAD with the true SSAD. I characterize SSAD shape by the number of sites occupied by a species and the maximum likelihood estimate of the negative binomial clustering parameter $k$. The results of this simulation are discussed in the main text.

```
simNBPerm <- ssadSim(nsite, nspp, nthrd, function(n) {rlseries(n, b)},
                     function(n, mu) {rnbinom(n, oneK, mu = mu)}, nsim = nsim)

simPoPerm <- ssadSim(nsite, nspp, nthrd, function(n) {rlseries(n, b)},
                     function(n, mu) {rpois(n, lambda = mu)}, nsim = nsim)
```

#### S6 Exploring the independent swap null model algorithm

As discussed in the main text, a null model algorithm that constrains the shape of the SSAD for each species might be robust to statistical artifacts such as spatial clustering of abundances. To evaluate if that is true I repeat the simulation experiment that produced community data with absolutely no real species associations while still maintaining SSAD and SAD shapes extracted from the data. In this new simulation experiment, however, I use the independent swap algorithm (Kembel *et al.*, 2010; Ulrich & Gotelli, 2010) instead of the fixed-fixed algorithm. The independent swap algorithm preserves the shape of the SSAD for each species

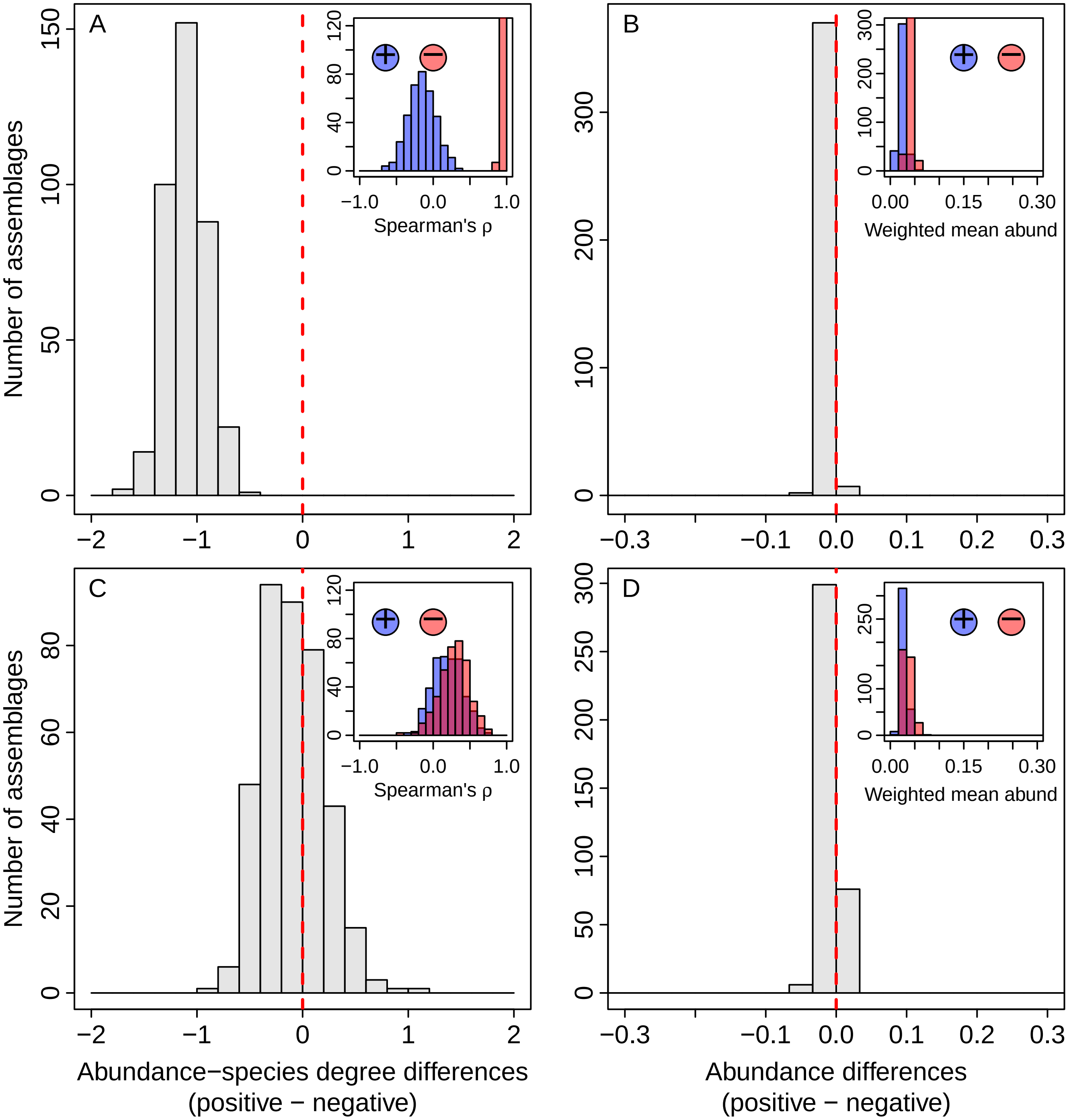


Supplementary Figure 7: Results from simulated abundances and association networks when only a single SAD shape and a single SSAD shape are used (as described in the text). Panel (A) shows correlations between abundance and centrality in positive or negative associations networks; panel (B) shows frequencies of different abundances in the two types of association networks.

while randomizing its occurrences. Despite the conservative nature of this null model algorithm it still detects spurious species association networks from simulated data, and still produces spurious relationships between abundances and positive or negative associations (Supp. Fig. 8). Supp. Fig. 8 shows the results of this simulation experiment and closely matches Supplementary Figure 2 from CEA, indicating that their results can be reproduced with data that contain no real species associations even when using the more conservative independent swap algorithm.

```
# re-set number of simulations to run
nsim <- round(1.25 * sum(!is.na(commStats$pos.wm)))

# explore independent swap algorithm instead of fixed-fixed algo
simIndSwapData <- indSwapTest(sadStats = sadStats, mcCores = nthrd,
                              ssadType = 'nbinom', kfun = kfun, nsim = nsim)
```

```
fig2bc(simIndSwapData, breaksRho = seq(-2, 2, by = 1/5),
       breaksWM = seq(-0.3, 0.3, by = 0.1/3))
```

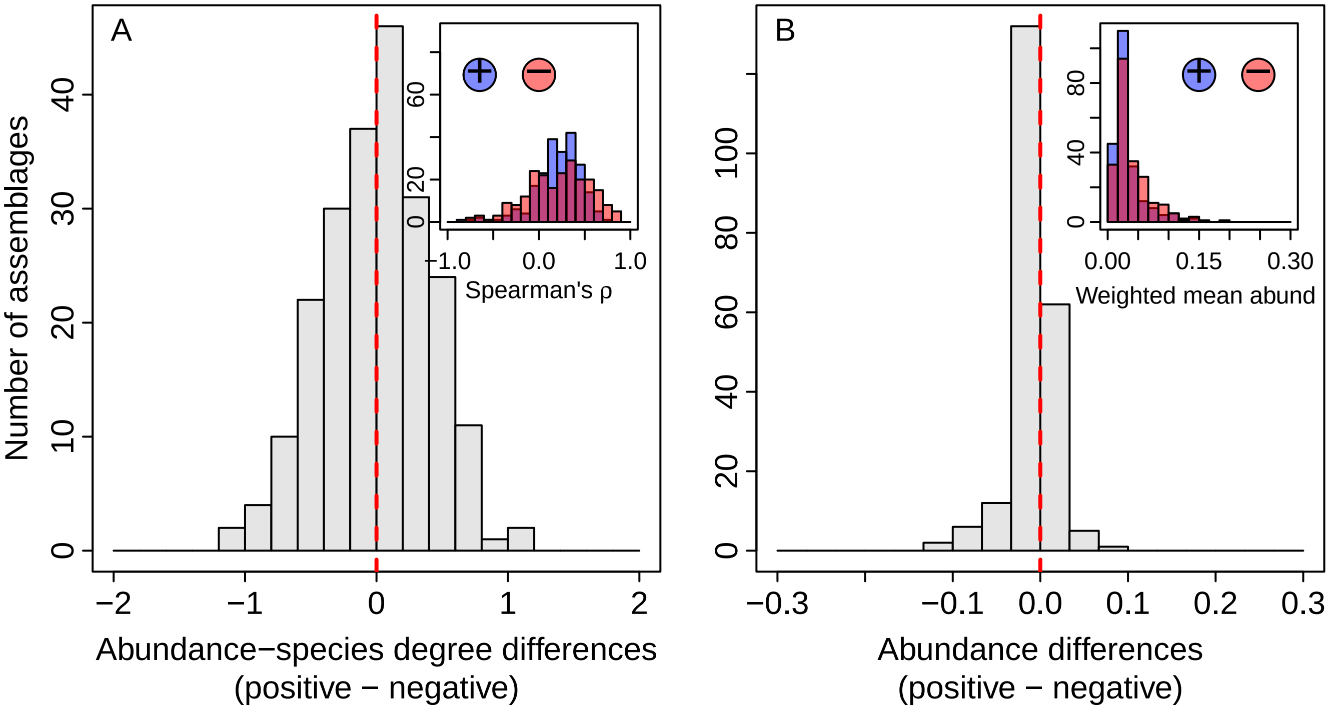


Supplementary Figure 8: Results from simulated abundances and association networks when the independent swap algorithm is used to infer species associations. Panel (A) shows correlations between abundance and centrality in positive or negative associations networks; panel (B) shows frequencies of different abundances in the two types of association networks.

#### S7 Mathematical exploration of different SSADs and Schoener similarity

Finally I very simply want to understand at a mathematical level why a spatially clustered versus spatially even SSAD would lead to these results. To explore this I consider a very simple example of two species and their Schoener similarity.

```r
N <- 5
Nrare <- 5
Ncomm <- 50
nsite <- 5

rarePair <- cbind(c(1, rep(0, nsite - 1)), c(Nrare, rep(0, nsite - 1)))
commPair <- cbind(c(Ncomm, rep(0, nsite - 1)), c(0, Ncomm, rep(0, nsite - 2)))
commPairMax <- matrix(rep(Ncomm / nsite, nsite * 2), ncol = 2)
```

First I consider two rare species, one with abundance 1, and the other with abundance 5. If I consider a dataset with 5 total sites, then the configuration that maximizes the Schoener similarity between these two species is

$$\begin{bmatrix} 1 & 5 \\ 0 & 0 \\ 0 & 0 \\ 0 & 0 \\ 0 & 0 \end{bmatrix}$$

I can calculate how likely this configuration is under a negative binomial versus a Poisson SSAD like this

```r
k <- 0.1
nbRareP <- prod(dnbinom(rarePair[, 2], k, mu = Nrare / nsite))
poRareP <- prod(dpois(rarePair[, 2], Nrare / nsite))
```

I see that the negative binomial SSAD is more likely to maximize the Schoener similarity compared to the Poisson, and thus compared to the null model rare species will appear to be aggregated with each other.

Conversely for the common species, in my simple example represented by two species both with abundance 50, I want to compare the probabilities of minimizing the Schoener similarity between them as derived from the negative binomial versus the Poisson SSAD. This occurs in any configuration such as this one where their total abundances fail to overlap

$$\begin{bmatrix} 50 & 0 \\ 0 & 50 \\ 0 & 0 \\ 0 & 0 \\ 0 & 0 \end{bmatrix}$$

I can calculate the probabilities of any such configuration like this

```r
nbCommP <- prod(dnbinom(commPair[, 1], k, mu = Ncomm / nsite)) * (nsite - 1) *
    prod(dnbinom(commPair[, 2], k, mu = Ncomm / nsite))

poCommP <- prod(dpois(commPair[, 1], Ncomm / nsite)) * (nsite - 1) *
    prod(dpois(commPair[, 2], Ncomm / nsite))
```

I can further compare this to a scenario that would maximize the Schoener similarity between common species:

$$\begin{bmatrix} 10 & 10 \\ 10 & 10 \\ 10 & 10 \\ 10 & 10 \\ 10 & 10 \end{bmatrix}$$

I calculate the probability of this configuration like

```r
nbCommPMax <- prod(dnbinom(as.vector(commPairMax), k, mu = Ncomm / nsite))
poCommPMax <- prod(dpois(as.vector(commPairMax), Ncomm / nsite))
```

The conclusions of these probability calculations are discussed in the main text.
